# Reconciling DNA replication and transcription in a hyphal organism: Spatial dynamics of transcription complexes in live *Streptomyces coelicolor* A3(2)

**DOI:** 10.1101/498634

**Authors:** Leena Nieminen, Paul A. Hoskisson

## Abstract

Reconciling transcription and DNA replication in the growing hyphae of the filamentous bacterium *Streptomyces* presents several physical constraints on growth due to their apically extending and branching, multigenomic cells and chromosome replication being independent of cell division. Using a GFP translational fusion to the β-subunit of RNA polymerase (*rpoC-egfp*), in its native chromosomal location, we observed growing *Streptomyces* hyphae using time-lapse microscopy throughout the lifecycle and under different growth conditions. The RpoC-eGFP fusion co-localised with DNA around 1.8 μm behind the extending tip, whereas replisomes localise around 4-5 μm behind the tip, indicating that at the growing tip, transcription and chromosome replication are to some degree spatially separated. Dual-labeled RpoC-egfp/DnaN-mCherry strains also indicate that there is limited co-localisation of transcription and chromosome replication at the extending hyphal tip. This likely facilitates the use of the same DNA molecule for active transcription and chromosome replication in growing cells, independent of cell division. This represents a novel, but hitherto unknown mechanism for reconciling two fundamental processes that utilise the same macromolecular template that allows for rapid growth without compromising chromosome replication in filamentous bacteria and may have implications for evolution of filamentous growth in microorganisms, where uncoupling of DNA replication from cell division is required.

## Introduction

The processes of transcription and chromosome replication both occupy the same cellular template and understanding how such conflicts are reconciled is fundamental to understanding the complexities of bacterial growth and the structure of the dynamic bacterial nucleoid^1,2,3^. In eukaryotes this problem is solved by segregating growth and replication in to separate stages within the cell cycle. In bacteria, this is not the case and spatial organisation of the nucloids is dependent on the growth habits and morphology of the specific bacterium^1^. Bacterial RNAP is highly sensitive to environmental cues and is subject to significant compaction and expansion forces due to the action of DNA-binding proteins, DNA supercoiling, macromolecular crowding, interaction with cytoskeletal proteins and transertion^4,5^ impacting on other cell processes such as DNA replication. *Streptomyces* are filamentous saprophytic bacteria that have a complex lifecycle, where a single unigenomic spore gives rise to a multi-compartment, multi-genomic vegetative hyphal mass that can forage for nutrients through tip extension. In response to nutrient limitation or stress, specialised multigenomic aerial hyphae are raised in to the air that form septa, resulting in the formation of a unigenomic compartment which completes development in to a mature spore^6,7^. This hyphal growth habit is remarkably similar to that of the filamentous fungi and represents an excellent example of how two groups of organisms have adapted to life in soil through convergent evolution. Several aspects of *Streptomyces* biology challenge our understanding of bacterial nucleoid structure/function and cell division, its links to chromosome replication and segregation and how this is reconciled with transcriptional activity. The large (8-10 Mbp) linear chromosome found in *Streptomyces*, appears to be largely uncondensed during vegetative growth^8^ but is highly ordered in terms of its structure and transcriptional activity^9^ and unlike the majority of bacteria it can be replicated independently of cell division^10^. *Streptomyces* are unusual amongst bacteria as many of the genes required for cell division are dispensable for vegetative growth such as *ftsZ, ftsQ*, and *mreB*, contrary to what is observed in unicellular bacteria^10–12^. The temporal and spatial location and activity of key cellular proteins and nucleoids in *Streptomyces* is likely to have significant implications for our understanding of growth and development in hyphal bacteria. It is known that chromosome replication does not occur at the apex of hyphal tips in Streptomyces^8,13,14^ yet it is asynchronous and non-uniform along extending hyphae ^8^ What is less well understood is whether there is any hierarchical organisation of transcription in growing *Streptomyces* hyphae. In unicellular bacteria transcriptional foci or patches occur in discrete locations in rapidly growing cells and are associated with the rRNA operons in bacterial chromosomes^15–18^. Recently we have begun to understand the evolutionary mechanisms that minimise these conflicts in unicellular bacteria such as genome organisation, avoidance of co-occupancy and recycling of stalled replisomes/RNA polymerase (RNAP) holoenzyme on DNA^2,3^. In *Streptomyces* however, the hyphal lifestyle represents a fundamental evolutionary problem, that is, to reconcile the issues of chromosome replication and transcription in tandem with the structural complications of the presence of linear chromosomes and that chromosome replication is independent of cell division. To attempt to understand this problem we made a translational fusion of *rpoC* with *egfp* in its native chromosomal location and studied the dynamics of transcription throughout the lifecycle of *Streptomyces* using time-lapse microscopy in live cells.

## Materials and Methods

### Bacterial strains, plasmids, growth conditions and conjugal transfer from *E. coli* to *Streptomyces*

The *S. coelicolor* strains and cosmids used in this study are summarised in Table 1. All strains were grown on mannitol and soya flour (MS) agar^19^, solid nutrient agar^20^ or minimal medium with mannitol^21^. Conjugation from the *E. coli* strain ET12567 (*dam*^−^ *dcm*^−^ *hsdS*) containing the driver plasmid pUZ8002, was used to bypass the methyl-specific restriction system of *S. coelicolor* ^21^.

**Table 1.** Strains and plasmids used in this study

| Strain or plasmid | Genotype/comments | Source or reference |
| --- | --- | --- |
| ***S. coelicolor* strains** |  |  |
| M145 | Prototrophic, SCP1<sup>-</sup> SCP2<sup>-</sup> | 21 |
| sLN301 | Prototrophic, SCP1<sup>-</sup> SCP2<sup>-</sup>; *rpoC-egfp* | This work. |
| M570 | $\Delta relA$ | 29 |
| sLN401 | $\Delta relA$; *rpoC-egfp* | This work. |
| DJ542 | M145 *dnaN-mCherry* - unmarked with antibiotic resistance | Jakimowicz, Unpublished |
| sLN501 | M145, *rpoC-egfp* fusion in a DJ542 background – dual GFP & mCherry fluorescence | This work. |
| **Cosmids** |  |  |
| D40A | SuperCos derived cosmid vector with a genomic fragment containing the *rpoC* gene. | 41 |
| pLN301 | Cosmid D40A with an in-frame *eGFP* fusion to the 3' end of *rpoC* gene | This work. |

### Construction of the RpoC-eGFP fusion strains

The *rpoC-egfp* fusion was created using ReDirect technology^22^ in its native chromosomal location. The *egfp-aac(3)IV-oriT* cassette was amplified using oligonucleotides containing 39 nucleotide homologous extensions to chromosomal sequence of the 3’ end of *rpoC* (SCO4655) and its adjacent flanking region (For - 5’-CCGCTGGAGGACTACGACTACGGTCCGTACAACCAGTACCTGCCGGGCCCGGGCTGCCGGGCCCGGAGGTGAGCAAGGGCGAGGAGCT-3’ and Rev - 5’-CTCGGGGTGACCGCCCTTCGGTCGTATCAAGCTGCCCGCTTCCGGGGATCCGTCGACC-3’) as used by Ruban-Osmialowska et al.,^8^ in cosmid D40A, creating cosmid pLN301 (*rpoC-egfp*). The cosmid, pLN301 was moved in to the non-methylating *E. coli* strain ET12567/pUZ8002 to facilitate conjugation in to *S. coelicolor*, creating strain sLN301 (M145; *rpoC-egfp)* and was confirmed by sequencing and Southern hybridization (data not shown). Cosmid pLN301 was also moved in to the *relA* deletion strain M570 (*hyg* resistant) and mutant strains were selected on hygromycin and apramycin resistance, kanamycin sensitivity, creating sLN401. In addition pLN301 was conjugated in to DJ542, an unmarked *dnaN-mCherry* fusion. Strains were confirmed by sequencing and Southern hybridization (data not shown).

Using fluorescent microscopy and a previously established time-lapse fluorescent microscopy procedure^23^ we monitored RpoC-eGFP as a reporter of RNAP spatial and temporal dynamics under a range of conditions (see Results). Nucleic acid staining was achieved using SYTO42 (Life Technologies Corp.) and membranes were stained using FM4-64 (Life Technologies Corp.) according to the manufacturers instructions. Images were captured using a Nikon TE2000S inverted fluorescence microscopy. Exposure times were 20 ms for phase-contrast and 100 ms for fluorescence imaging. Images were analysed using IPLab scientific imaging software version 3.7 (Scanalytics, Inc., Rockville, USA). Statistical analysis was performed using Microsoft Office Excel software.

## Results and Discussion

### RpoC-eGFP patches show dynamic localisation throughout the lifecycle of *Streptomyces coelicolor*

To determine the location and dynamics of RNAP during the complex lifecycle of *S. coelicolor* we constructed a fusion of eGFP to the β’ subunit of RNAP core enzyme (SCO4655 ^15–18,24^). The *rpoC-egfp* fusion strain (sLN301) was found to sporulate normally and to grow at the same apical extension rate as the wild-type strain, enabling us to conclude that the fusion protein was functional (Fig. 1). We observed the location of RNAP throughout the lifecycle of *S. coelicolor* (Fig. 1) by monitoring RpoC-eGFP localization in combination with fluorescence stains for nucleic acids (SYTO42) and cell membranes (FM4-64).

**Fig. 1.**
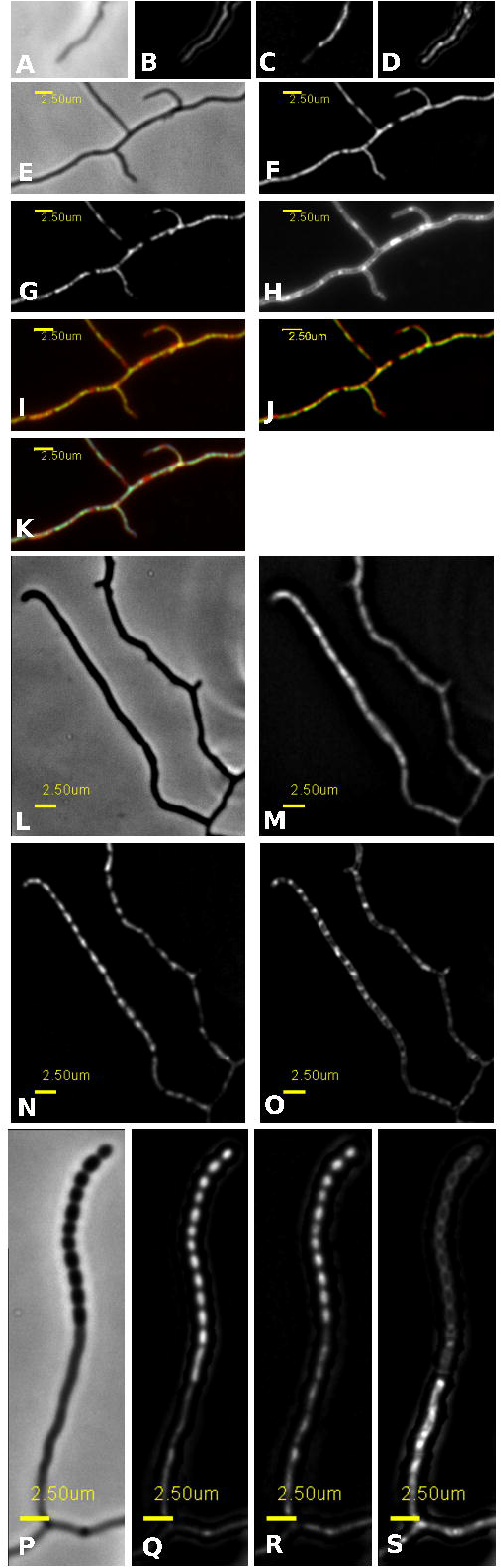
RpoC-eGFP patches show dynamic localisation throughout the lifecycle of *Streptomyces coelicolor*. Representative images of a germinating spore in phase contrast **(A)**, germinating spore stained with SYTO 42 (DNA staining; **B**), RpoC-eGFP localisation in a germinating spore **(C)**, germinating spore stained with FM4-64 (membrane stain; **D**). Representative images of vegetative hyphae in phase contrast **(E)**, vegetative hyphae stained with SYTO 42 (DNA staining; **F**), RpoC-eGFP localisation in a vegetative hypha **(G)**, vegetative hyphae stained with FM4-64 (membrane stain; **H**), a multiprobe image (RNAP-eGFP in green and FM4-64 in red; **I**), a multiprobe image (RpoC-eGFP in green and SYTO 42 in red; **J**), a multiprobe image (RpoC-eGFP, FM4-64 & SYT042; **K**). Representative images of aerial hyphae in phase contrast **(L)**, aerial hyphae stained with SYTO 42 (DNA staining; **M**), RpoC-eGFP localisation in an aerial hypha **(N)**, aerial hypha stained with FM4-64 (membrane stain; **O**). Representative images of a spore chain in phase contrast **(P)**, a spore chain stained with SYTO 42 (DNA staining; **Q**), RpoC-eGFP localisation in a spore chain **(R)**, a spore chain stained with FM4-64 (membrane stain; **S**).

RNAP was distributed throughout the apically extending germ tubes of sLN301 (*rpoC-egfp*) and co-localised with nucleic acids stained with SYTO42 (Fig. 1 A-D). Localisation of RNAP and nucleic acids was found to be in close proximity to the extending hyphal tip (< 1 μm). As the extending hyphae mature, the distance between RNAP and the apically extending tip increases. These branching vegetative hyphae exhibit distinct nucleic acid (nucleoid) patches that co-localise with RNAP in distinct areas within the hyphae (Fig. 1. E-K; See below also). Moreover the distance from the tip to the first RNAP patch appears to be around 2 μm throughout the vegetative mycelium (1.8 μm ±0.3 μm; n=29), suggesting that transcription is spatially constrained at the extending tip as observed in other hyphae.

Examining the distribution RNAP during the growth of aerial hyphae indicated that RNAP and nucleic acids were distributed throughout the extending aerial hyphae without showing the discrete pattern behind the extending tip observed in vegetative hyphae (Fig. 1, L-O). This may represent the requirement for complete distribution of transcriptional activity throughout the aerial hyphae for the maturation of spore chains. Examination of mature spore chains show that RNAP co-localised with the condensed and segregated nucleoids within the septated spore chains (Fig. 1, P-S).

### RNAP tracks behind the extending hyphal tip

To characterize the dynamics of RNAP in extending hyphae time-lapse images of *S. coelicolor* sLN301 (*rpoC-egfp*) were generated as phase-contrast images merged with GFP images (FITC filter) every 30 minutes during growth on minimal medium plus mannitol as a carbon source. RpoC-eGFP was observed in discrete patches and tracked behind the extending hyphal tip (Fig. 2A) at a mean distance of 1.8 μm (±0.3 μm; n=29) with the dimensions of the patches being 2.5 μm (+/− 1.6 μm; n=116). The emerging branches on the vegetative hyphae also showed the same distribution pattern of RpoC-eGFP patches as the extending primary hyphae. There appears to be some variation in the intensity of the RNAP-eGFP patches within the hyphae, although no obvious pattern could be determined, it may be that this variation is due to the differences in expression levels of various regions in the genome, such as the rRNA operons^15–18^.

**Fig. 2.**
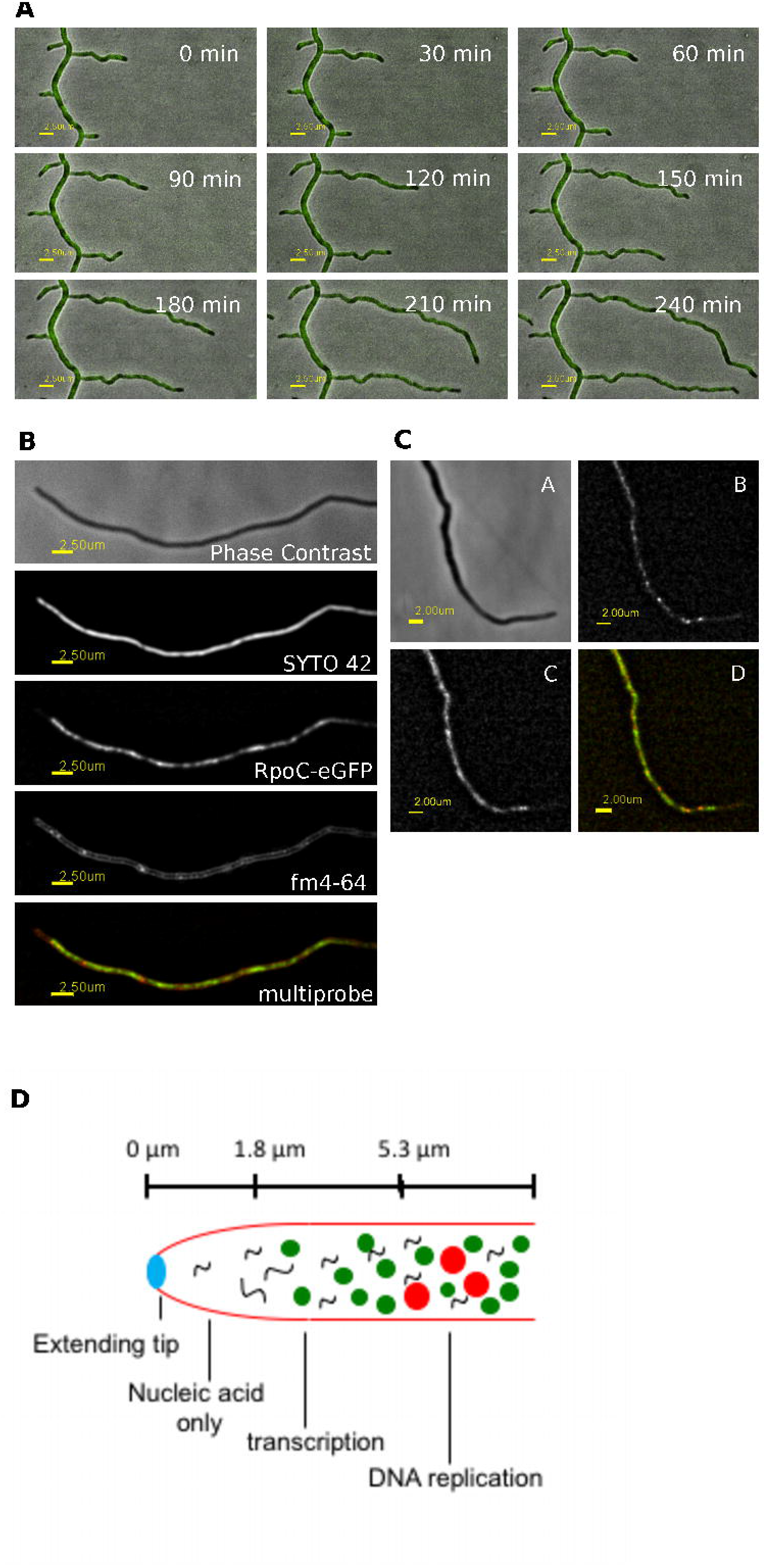
RpoC-eGFP patches track behind the extending hyphal tip. **(A)** Time-lapse images of growing *S. coelicolor* hyphae (LN301; *rpoC-egfp*) showing the absence of RNAP-eGFP patches at the tip of extending vegetative hyphae. (See also Supplementary video 1 - http://dx.doi.org/10.6084/m9.figshare.1181785) **B: RpoC-eGFP patches co-localise with DNA, but not at the hyphal tip.** Representative images of a vegetative hypha in phase contrast, stained with SYTO 42 (DNA staining), RNAP-eGFP, FM4-64 (membrane stain) and a multiprobe image (RNAP-eGFP in green and FM4-64 in red). **C: The majority of RpoC-eGFP patches do not co-localise with DnaN-mCherry at the hyphal tip, but do co-localise behind the tip.** Representative images of a vegetative hypha in phase contrast **(A)**, DnaN-mCherry **(B)** RNAP-eGFP **(C)** and a multiprobe image **(D)** of RNAP-eGFP (green) and DnaN-mCherry (Red). **D**: Schematic representation of a hyphal tip (polarisome), indicating the locations of nucleic acids, transcription (this work) and replisome location^14–18^ suggesting there is a spatial separation of transcription and chromosome replication at the hyphal tip.

### RNAP patches co-localise with DNA but not at the hyphal tip

Examining vegetative hyphae by phase contrast, RNAP-eGFP (FITC filter) and fluorescent staining of nucleic acids (SYTO42) and membranes (FM4-64) it can be seen that RNAP patches clearly co-localize with DNA (Fig. 2B). However nucleic acids stained by SYTO 42 extends to the hyphal tip, whereas RNAP-eGFP was never observed at the tip of extending hyphae. When compared to the patches for replisomes, measured by Wolanski et al.,^14^ at 5.3 μm (± 2.0 μm) behind the hyphal tip, the RNAP-eGFP patches were found located at a mean of 1.8 μm behind the extending tip suggesting there is a spatial separation of transcription and DNA replication at the hyphal tip. These data, obtained from single tagged strains, suggest that one or more chromosomes are actively transcribing at the extending tip, yet active replication occurs behind this. To further examine this spatial separation hypothesis, a double fluorescent strain *dnaN-mCherry/rpoC-egfp* (sLN501) was constructed. In sLN501 (*dnaN-mCherry/rpoC-egfp*) RNAP patches were observed to lag behind the tip, as previously observed and DnaN-mCherry tagged replication factories were located distal to these. Discrete RpoC-eGFP patches, un-associated with DnaN-mCherry were observed proximal to the extending tip (Fig 2C), further supporting our hypothesis of spatially separated transcription and DNA replication at the apical tip of extending *Streptomyces* hyphae. These data suggest there is a hierarchy of chromosome occupancy at the tip of extending hyphae that is summarized in our model (Fig. 2D). Whilst the molecular mechanism underpinning this spatial constraint is currently unknown, it is thought that avoiding co-occupancy of the DNA template occurs, at least to some extent, in eukaryotes^25^. The unusual combination of linear chromosomes and apical growth in *Streptomyces*, coupled with DNA replication being independent of cell division and chromosome segregation, suggests that this mechanism may have evolved to allow active transcription at the actively growing tips, independent of DNA replication and cell division. This is consistent with the replisome trafficking data of Wolanski et al.,^14^ and intriguingly could involve the pleiotrophic regulator AdpA, which has recently been shown to control chromosome replication through competition with DnaA at *oriC*^26^.

### RNAP shows *relA-dependant* pausing during nitrogen starvation

To investigate how environmental cues may affect RNAP dynamics in *S. coelicolor* we examined the effect of the stringent response on RNAP localisation. The highly phosphorylated guanosine nucleotide ppGpp is known to mediate growth rate dependent gene expression in bacteria through direct interaction with RNAP during the stringent response^27,28^. In *Streptomyces*, ppGpp is synthesised by RelA, and has previously been shown to influence control over antibiotic production and morphological development in response to nutrient limitation^29–31^, however, what is not known is how RelA influences the dynamics of RNAP within *Streptomyces* cells in response to nutrient downshift. To test this, we grew *S. coelicolor* sLN301 (WT *rpoC-egfp*) and sLN401 (*ΔrelA rpoC-egfp*) on cellophane discs placed upon on solid nutrient agar (Rich medium, amino acid/peptide based nitrogen source). Once cells were growing exponentially, cellophane squares were removed and applied to minimal medium containing sodium nitrate as the sole nitrogen source (30 mM;^32^) to induce nitrogen-starvation and the stringent response. Following nitrogen downshift, the dynamics of RNAP patches was followed (Fig. 3), in strain sLN301 (WT *rpoC-egfp*) cell growth paused and RpoC-eGFP patches remained static, presumably during the stringent response and the synthesis of ppGpp by RelA. After 60 mins mycelial growth resumed, but from new branch points in the mycelium and following 120 mins, apical growth was within the normal distribution range of RpoC-eGFP patches. The resumption of growth via branching is intriguing and may involve the serine/threonine protein kinase, AfsK. It is known that branching is affected by environmental conditions^33^ and that AfsK plays a role in the onset of secondary metabolism and sporulation, both nutrient dependent processes^34–36^. Recently it has been shown that AfsK co-localizes and directly regulates DivIVA in *Streptomyces*^36,37^. Induction of AfsK results in branching and it is believed that phosphorylation of DivIVA results in disassembly of the apical polarisome and the assembly of new growth patches at branch points. Interestingly this may represent a mechanism of altering growth habit in response to nutrient limitation, increasing the nutrient foraging ability of bacterial colonies. Repeating the experiment with sLN401 (*ΔrelA rpoC-egfp*) resulted in no cessation of growth and no increased branching following nitrogen-downshift. Intriguingly this suggests a role for the stringent response in reprogramming the growth habit (apical growth and branching) of *Streptomyces* in response to nitrogen-downshift, however neither AfsK or DivIVA were identified as direct targets in a microarray study of a *ΔrelA* mutant and a ppGpp inducible strain^38^, suggesting there is an as yet unknown mechanism integrating these signals.

**Fig. 3.**
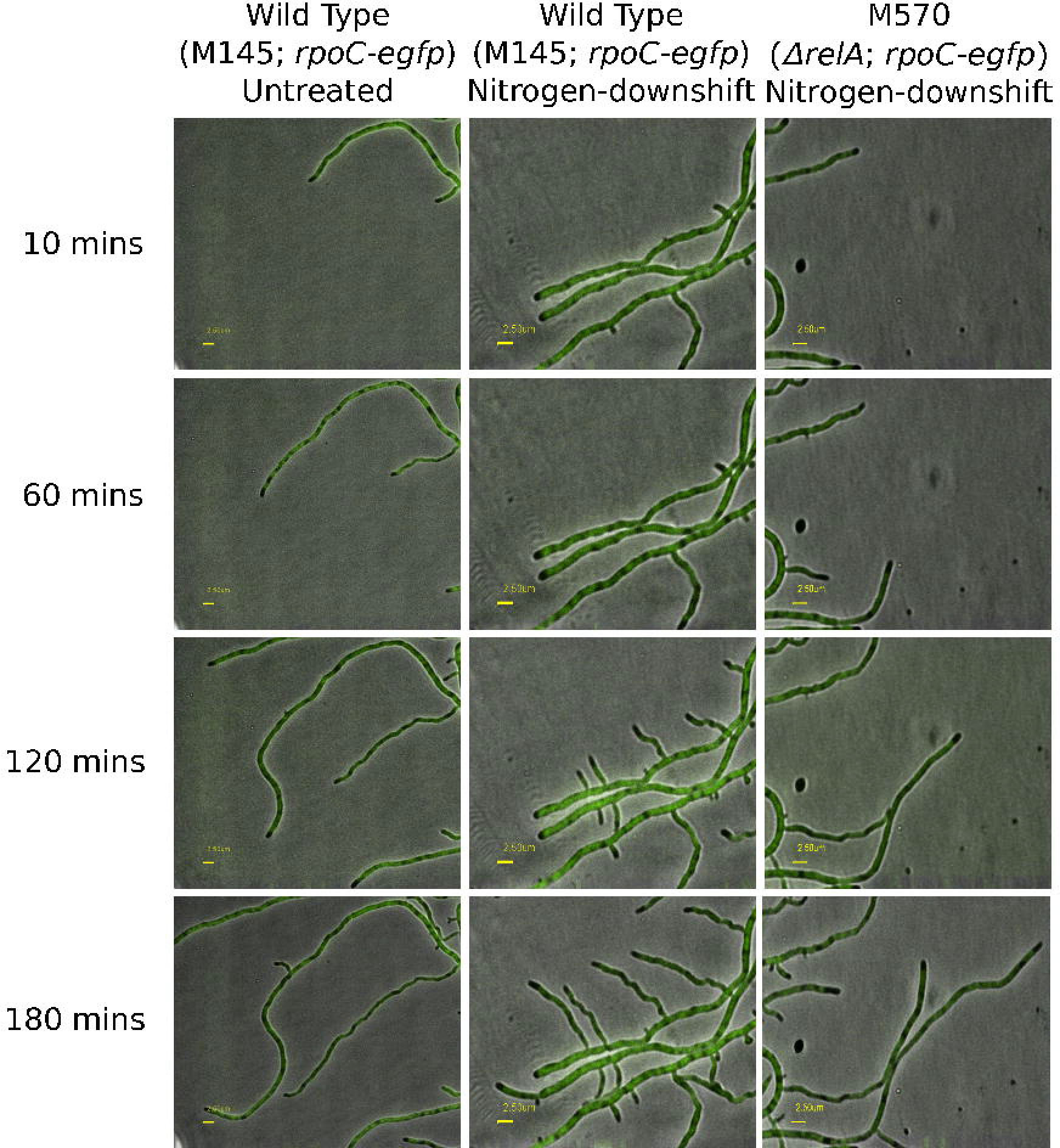
RpoC-eGFP patches in Wild-Type *S. coeliocolor* exhibit pauses following nitrogen-downshift when compared to a *ΔrelA* mutant. Time-lapse images of growing *S. coelicolor* hyphae (sLN301; *rpoC-egfp*) in nitrogen rich (nutrient agar) medium over 180 min. (See also Supplementary video 2 - http://dx.doi.org/10.6084/m9.figshare.1181781). Time-lapse images of growing *S. coelicolor* hyphae (sLN301; *rpoC-egfp*) following nitrogen downshift over 180 min. (See also Supplementary video 3 - http://dx.doi.org/10.6084/m9.figshare.1181780). Time-lapse images of growing M570 *S. coelicolor* hyphae (*ΔrelA; rpoC-egfp*) following nitrogen downshift over 180 min. (See also Supplementary video 4 - http://dx.doi.org/10.6084/m9.figshare.1181782)

### Disruption of transcription or translation results in altered RNAP dynamics in hyphae

To further understand the dynamics of RNAP in live *S. coelicolor* hyphae, we used antibiotic rifampicin to inhibit transcription and chloramphenicol to inhibit translation. *S. coelicolor* sLN301 (WT *rpoC-egfp*) was grown in the absence of each antibiotic on cellophane, once cells were growing exponentially, cellophane squares were removed and applied to the same medium containing 50 % of the minimum inhibitory concentrations (MIC) of each antibiotic (Fig. 4). Treatment of *S. coelicolor* sLN301 (WT *rpoC-egfp*) with rifampicin resulted in no cessation of the apical extension rate of hyphae, however RpoC-eGFP patches became dispersed, consistent with disassociation of RNAP from the nucleoid (Fig. 4); resulting in an overall increase in the size of fluorescent patches from 2.5 μm (± 1.5 μm; n=54) in untreated to 4.3 μm (± 3.0 μm; n=30). Rifampicin inhibits initiation and re-initiation of transcription through targeting β-subunit of RNAP core enzyme and this dispersal of RNAP patches following rifampicin treatment has also been observed in *Escherichia coli*^17^. Treatment of sLN301 (WT *rpoC-egfp*) with chloramphenicol resulted in a cessation of apical extension over a 120 min period and condensation of the RpoC-eGFP patches (Fig. 4), which is consistent with observations in other organisms^39^. The RpoC-eGFP patches also move away from the apical tip following treatment 2.0 μm (± 0.4 μm; n=14) in untreated to 4.5 μm (± 2.5 μm; n=15). Moreover, it has also been shown that active transcription is required for such compaction^17^ suggesting that the compaction observed in *S. coelicolor* indicates that transcriptional activity is occurring in these patches and that active transcription is not occurring at the tip as shown above (Fig.1). The coupling of transcription and translation in bacteria has potentially profound effects on the structure of the nucleoid^17^, the two antibiotics used in this study both inhibit translation, but in different ways; chloramphenicol directly inhibits translation, but does not prevent transcription, yet rifampicin inhibits transcription and due to the coupling of these processes in bacteria it also inhibits translation^17^. It has also been shown that transcriptional activity is adjusted in bacteria to meet the translational needs of cells under various growth conditions ^40^ suggesting that mechanisms to reconcile potentially conflicting key cellular processes such as transcription, translation and DNA replication can help reduce the extreme effects such process can have on growth and nucleoid structure.

**Fig. 4.**
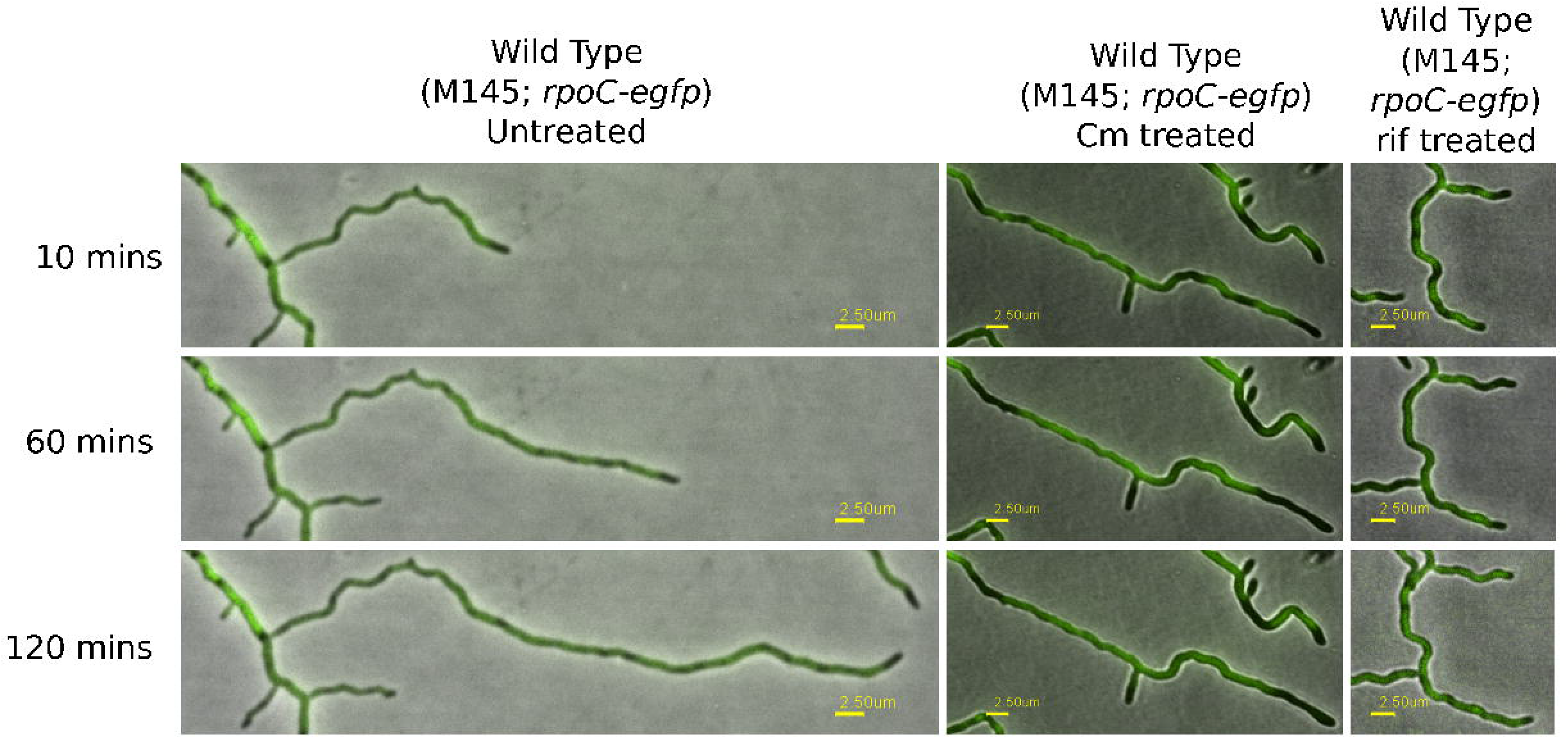
RpoC-eGFP patches exhibit altered dynamics following inhibition of either transcription or translation. Time-lapse images of growing *S. coelicolor* hyphae (sLN301; *rpoC-egfp*) without any antibiotic treatment. Time-lapse images of growing *S. coelicolor* hyphae (sLN301; *rpoC-egfp)* following treatment with chloramphenicol (Cm; 13 mg ml^−1^). See also Supplementary video 5 - http://dx.doi.org/10.6084/m9.figshare.1181783. Time-lapse images of growing *S. coelicolor* hyphae (sLN301; *rpoC-egfp*) following treatment with rifampicin (rif; 32 mg ml^−1^). See also Supplementary video 6 - http://dx.doi.org/10.6084/m9.figshare.1181784.

## Summary

The tip growth habit of *Streptomyces* challenges our understanding of how transcription and replisome occupancy of the same template in bacteria can occur. One way to resolve this is to spatially separate the two processes. Intriguingly, eukaryotic organisms temporally separate key cellular processes such as growth and replication. The data presented here suggest that the tip of the actively growing *Streptomyces* hyphae spatial separates DNA replication and transcription. In these rapidly extending areas of the mycelium, transcription and replication on the same template may lead to collisions, and separating these transcribing nucleoids from replicating nucleoids offers an attractive means to achieving this. Whilst the mechanism of this spatial separation is currently unknown, spatial or temporal separations of conflicting processes is an attractive mechanism to maximise apical growth with minimal conflict between transcription and DNA replication. This may be especially important for soil organisms such as *Streptomyces* or fungi that, through convergent evolution, exhibit similar apical growth habits in a resource-limited ecological niche.

## Acknowledgements

We would like to thank Dr Dagmara Jakimowicz, University of Wroclaw, Poland for the gift of the dnaN-mCherry strain and helpful comments on the manuscript. We would also like to thank Prof. Mervyn Bibb FRS and Dr Andrew Hesketh of the John Innes Centre for the gift of the *relA* mutant. We would like to thank Dr Paul R Herron, University of Strathclyde, for microscopy assistance and discussions.

## Supplementary data

**Supp Video 1: RpoC-eGFP patches tracking behind the extending hyphal tip.** Video of growing *S. coelicolor* hyphae (LN301; *rpoC-egfp*) showing the absence of RNAP-eGFP patches at the tip of extending vegetative hyphae. Images taken at 10 min intervals and converted to video using IPLab scientific imaging software version 3.7 (Scanalytics, Inc., Rockville, USA). http://dx.doi.org/10.6084/m9.figshare.1181785

**Supp Video 2: RpoC-eGFP patches in Wild-Type *S. coelicolor*.** Video of growing *S. coelicolor* hyphae (sLN301; *rpoC-egfp*) in nitrogen rich (nutrient agar) medium over 180 min. Images taken at 10 min intervals and converted to video using IPLab scientific imaging software version 3.7 (Scanalytics, Inc., Rockville, USA). http://dx.doi.org/10.6084/m9.figshare.1181781

**Supp Video 3: RpoC-eGFP patches in Wild-Type *S. coeliocolor* exhibit pauses following nitrogen-downshift.** Video of growing WT *S. coelicolor* hyphae (sLN301; *rpoC-egfp*) following nitrogen downshift over 180 min. Images taken at 10 min intervals and converted to video using IPLab scientific imaging software version 3.7 (Scanalytics, Inc., Rockville, USA). http://dx.doi.org/10.6084/m9.figshare.1181780

**Supp Video 4: RpoC-eGFP patches in a *ΔrelA* mutant of *S. coeliocolor* exhibit pauses following nitrogen-downshift.** Video of growing *S. coelicolor* hyphae (Δ*relA; rpoC-egfp*) following nitrogen downshift over 180 min. Images taken at 10 min intervals and converted to video using IPLab scientific imaging software version 3.7 (Scanalytics, Inc., Rockville, USA). http://dx.doi.org/10.6084/m9.figshare.1181782

**Supp Video 5: RpoC-eGFP patches exhibit altered dynamics following inhibition of translation.** Video of growing *S. coelicolor* hyphae (sLN301; *rpoC-egfp*) following treatment with chloramphenicol (Cm; 13 mg ml^−1^). Images taken at 10 min intervals and converted to video using IPLab scientific imaging software version 3.7 (Scanalytics, Inc., Rockville, USA). http://dx.doi.org/10.6084/m9.figshare.1181783

**Supp Video 6: RpoC-eGFP patches exhibit altered dynamics following inhibition of transcription.** Video of growing *S. coelicolor* hyphae (sLN301; *rpoC-egfp*) following treatment with rifampicin (rif; 32 mg ml^−1^). Images taken at 10 min intervals and converted to video using IPLab scientific imaging software version 3.7 (Scanalytics, Inc., Rockville, USA). http://dx.doi.org/10.6084/m9.figshare.1181784

## References

1. Kois-Ostrowska, A. et al. Unique Function of the Bacterial Chromosome Segregation Machinery in Apically Growing *Streptomyces* - Targeting the Chromosome to New Hyphal Tubes and its Anchorage at the Tips. PLoS Genet 1–25 (2016).

2. Merrikh, H., Zhang, Y., Grossman, A. D. & Wang, J. D. Replication–transcription conflicts in bacteria. Nat Rev Micro 10, 449–458 (2012).

3. McGlynn, P., Savery, N. J. & Dillingham, M. S. The conflict between DNA replication and transcription. Molecular Microbiology 85, 12–20 (2012).

4. Woldringh, C. L. The role of co-transcriptional translation and protein translocation (transertion) in bacterial chromosome segregation. Molecular Microbiology 45, 17–29 (2002).

5. Cabrera, J. E., Cagliero, C., Quan, S., Squires, C. L. & Jin, D. J. Active Transcription of rRNA Operons Condenses the Nucleoid in *Escherichia coli*: Examining the Effect of Transcription on Nucleoid Structure in the Absence of Transertion. J. Bacteriol. 191, 4180–4185 (2009).

6. Flärdh, K. & Buttner, M. J. Streptomyces morphogenetics: dissecting differentiation in a filamentous bacterium. Nat Rev Micro 7, 36–49 (2009).

7. Hoskisson, P. A., Rigali, S., Fowler, K., Findlay, K. C. & Buttner, M. J. DevA, a GntR-like transcriptional regulator required for development in *Streptomyces coelicolor*. J. Bacteriol. 188, 5014–5023 (2006).

8. Ruban-Osmialowska, B., Jakimowicz, D., Smulczyk-Krawczyszyn, A., Chater, K. F. & Zakrzewska-Czerwinska, J. Replisome Localization in Vegetative and Aerial Hyphae of *Streptomyces coelicolor*. J. Bacteriol. 188, 7311–7316 (2006).

9. McArthur, M. & Bibb, M. In vivo DNase I sensitivity of the *Streptomyces coelicolor* chromosome correlates with gene expression: implications for bacterial chromosome structure. Nucleic acids research 34, 5395–5401 (2006).

10. McCormick, J. R. Cell division is dispensable but not irrelevant in *Streptomyces*. Curr. Opin. Microbiol. 12, 689–698 (2009).

11. McCormick, J. R., McCormick, J. R., Losick, R. & Losick, R. Cell division gene ftsQ is required for efficient sporulation but not growth and viability in *Streptomyces coelicolor* A3(2). 178, 5295–5301 (1996).

12. Mazza, P. et al. MreB of *Streptomyces coelicolor* is not essential for vegetative growth but is required for the integrity of aerial hyphae and spores. Molecular Microbiology 60, 838–852 (2006).

13. Yang, M. C. & Losick, R. Cytological evidence for association of the ends of the linear chromosome in *Streptomyces coelicolor*. J. Bacteriol. 183, 5180–5186 (2001).

14. Wolanski, M. et al. Replisome Trafficking in Growing Vegetative Hyphae of *Streptomyces coelicolor* A3(2). J. Bacteriol. 193, 1273–1275 (2011).

15. Lewis, P. J. Bacterial subcellular architecture: recent advances and future prospects. Molecular Microbiology 54, 1135–1150 (2004).

16. Lewis, P. J., Doherty, G. P. & Clarke, J. Transcription factor dynamics. Microbiology 154, 1837–1844 (2008).

17. Cabrera, J. E. & Jin, D. J. The distribution of RNA polymerase in *Escherichia coli* is dynamic and sensitive to environmental cues. Molecular Microbiology 50, 1493–1505 (2003).

18. Migocki, M. D., Lewis, P. J., Wake, R. G. & Harry, E. J. The midcell replication factory in *Bacillus subtilis* is highly mobile: implications for coordinating chromosome replication with other cell cycle events. Molecular Microbiology 54, 452–463 (2004).

19. Hobbs, G., Frazer, C., Gardner, D. J., Cullum, J. & Oliver, S. Dispersed growth of *Streptomyces* in liquid culture. Applied Microbiology and Biotechnology 31, (1989).

20. Hoskisson, P. A., Hobbs, G. & Sharples, G. P. Response of *Micromonospora echinospora* (NCIMB 12744) spores to heat treatment with evidence of a heat activation phenomenon. Letters in Applied Microbiology 30, 114–117 (2000).

21. Kieser, T., Bibb, M. J., Buttner, M. J., Chater, K. F. & Hopwood, D. A. Practical Streptomyces Genetics. (John Innes Foundation, 2000).

22. Gust, B., Challis, G. L., Fowler, K., Kieser, T. & Chater, K. F. PCR-targeted Streptomyces gene replacement identifies a protein domain needed for biosynthesis of the sesquiterpene soil odor geosmin. Proceedings of the National Academy of Sciences 100, 1541–1546 (2003).

23. Jyothikumar, V., Tilley, E. J., Wali, R. & Herron, P. R. Time-lapse microscopy of *Streptomyces coelicolor* growth and sporulation. Applied and Environmental Microbiology 74, 6774–6781 (2008).

24. Bentley, S. D. et al. Complete genome sequence of the model actinomycete *Streptomyces coelicolor* A3(2). Nature 417, 141–147 (2002).

25. Sutherland, H. & Bickmore, W. A. Transcription factories: gene expression in unions? Nature Reviews Genetics 10, 457–466 (2009).

26. Wolanski, M., Jakimowicz, D. & Zakrzewska-Czerwinska, J. AdpA, key regulator for morphological differentiation regulates bacterial chromosome replication. Open Biology 2, 120097–120097 (2012).

27. Vrentas, C. E. et al. Still Looking for the Magic Spot: The Crystallographically Defined Binding Site for ppGpp on RNA Polymerase Is Unlikely to Be Responsible for rRNA Transcription Regulation. Journal of Molecular Biology 377, 551–564 (2008).

28. Dalebroux, Z. D. & Swanson, M. S. ppGpp: magic beyond RNA polymerase. Nat Rev Micro 10, 203–212 (2012).

29. Chakraburtty, R. & Bibb, M. The ppGpp synthetase gene (relA) of *Streptomyces coelicolor* A3(2) plays a conditional role in antibiotic production and morphological differentiation. J. Bacteriol. 179, 5854–5861 (1997).

30. Hesketh, A., Sun, J. & Bibb, M. Induction of ppGpp synthesis in *Streptomyces coelicolor* A3(2) grown under conditions of nutritional sufficiency elicits actII-ORF4 transcription and actinorhodin biosynthesis. Molecular Microbiology 39, 136–144 (2001).

31. Sun, J. H., Hesketh, A. & Bibb, M. Functional Analysis of relA and rshA, two relA/spoT homologues of *Streptomyces coelicolor* A3(2). J. Bacteriol. 183, 3488–3498 (2001).

32. Karandikar, A., Sharples, G. P. & Hobbs, G. Differentiation of *Streptomyces coelicolor* A3 (2) under nitrate-limited conditions. Microbiology 143, 3581–3590 (1997).

33. Nieminen, L., Webb, S., Smith, M. C. M. & Hoskisson, P. A. A flexible mathematical model platform for studying branching networks: experimentally validated using the model actinomycete, *Streptomyces coelicolor*. PLoS ONE 8, e54316 (2013).

34. Ueda, K., Umeyama, T., Beppu, T. & Horinouchi, S. The aerial mycelium-defective phenotype of Streptomyces griseus resulting from A-factor deficiency is suppressed by a Ser/Thr kinase of *S. coelicolor* A3(2). Gene 169, 91–95 (1996).

35. Umeyama, T., Lee, P. C., Ueda, K. & Horinouchi, S. An AfsK/AfsR system involved in the response of aerial mycelium formation to glucose in *Streptomyces griseus*. Microbiology 145, 2281–2292 (1999).

36. Hempel, A. M. et al. The Ser/Thr protein kinase AfsK regulates polar growth and hyphal branching in the filamentous bacteria *Streptomyces*. Proc. Natl. Acad. Sci. U.S.A. 109, E2371–9 (2012).

37. Flärdh, K., Richards, D. M., Hempel, A. M., Howard, M. & Buttner, M. J. Regulation of apical growth and hyphal branching in *Streptomyces*. Curr. Opin. Microbiol. 15, 737–743 (2012).

38. Hesketh, A., Chen, W. J., Ryding, J., Chang, S. & Bibb, M. The global role of ppGpp synthesis in morphological differentiation and antibiotic production in *Streptomyces coelicolor* A3(2). Genome Biol 8, R161 (2007).

39. van Helvoort, J. M., Kool, J. & Woldringh, C. L. Chloramphenicol causes fusion of separated nucleoids in *Escherichia coli* K-12 cells and filaments. J. Bacteriol. 178, 4289–4293 (1996).

40. Proshkin, S., Rahmouni, A. R., Mironov, A. & Nudler, E. Cooperation between translating ribosomes and RNA polymerase in transcription elongation. Science 328, 504–508 (2010).

41. Redenbach, M. et al. A set of ordered cosmids and a detailed genetic and physical map for the 8 Mb *Streptomyces coelicolor* A3(2) chromosome. Molecular Microbiology 21, 77–96 (1996).

